# Machine learning methods for predicting guide RNA effects in CRISPR epigenome editing experiments

**DOI:** 10.1101/2024.04.18.590188

**Authors:** Wancen Mu, Tianyou Luo, Alejandro Barrera, Lexi R. Bounds, Tyler S. Klann, Maria ter Weele, Julien Bryois, Gregory E. Crawford, Patrick F. Sullivan, Charles A. Gersbach, Michael I. Love, Yun Li

## Abstract

CRISPR epigenomic editing technologies enable functional interrogation of non-coding elements. However, current computational methods for guide RNA (gRNA) design do not effectively predict the power potential, molecular and cellular impact to optimize for efficient gRNAs, which are crucial for successful applications of these technologies.

We present “launch-dCas9” (machine LeArning based UNified CompreHensive framework for CRISPR-dCas9) to predict gRNA impact from multiple perspectives, including cell fitness, wild-type abundance (gauging power potential), and gene expression in single cells. Our launch-dCas9, built and evaluated using experiments involving >1 million gRNAs targeted across the human genome, demonstrates relatively high prediction accuracy (AUC up to 0.81) and generalizes across cell lines. Method-prioritized top gRNA(s) are 4.6-fold more likely to exert effects, compared to other gRNAs in the same cis-regulatory region. Furthermore, launch-dCas9 identifies the most critical sequence-related features and functional annotations from >40 features considered. Our results establish launch-dCas9 as a promising approach to design gRNAs for CRISPR epigenomic experiments.

## Introduction

CRISPRi (Clustered Regularly Interspaced Short Palindromic Repeats interference) is a modified version of CRISPR that enables perturbation of noncoding putative cis-regulatory elements (CREs) in order to experimentally assess their functional impact. CRISPRi has become an increasingly popular tool for identifying functional regulatory elements, allowing researchers to quantify the effects on gene expression and control of various biological processes.^1–3^ While it is still infeasible and unnecessary to perform CRISPRi experiments using all potential gRNAs for CREs of interest across various relevant biological systems, developing prediction models that use existing data to predict the likely experimental outcomes for any given gRNA is crucial for optimizing the design and execution of new experiments. Such computational prediction models will facilitate the identification of promising gRNA targets for CREs that influence various molecular or cellular outcomes, including cell fitness (resulting from the balance of cell survival and proliferation), gRNA abundance in wild-type cells (when introducing the gRNA alone), and expression of nearby genes. Moreover, prediction models can facilitate the discovery of new and effective therapeutic targets impacting disease-relevant molecular profiles, be it cell fitness or expression of disease causing genes. Using gRNAs that more effectively target distal regulatory elements contributing to diseases enhances our potential to develop more effective treatments.

Published prediction models for optimal gRNA design are mainly developed for data from genetic (i.e., CRISPR) instead of epigenetic (CRISPRi or CRISPRa) perturbation experiments. The models built from CRISPR data focus primarily on predicting the efficiency of cutting the target sequence by Cas9 nuclease and off-target activity, commonly measured by indel frequency^4–9^. To our knowledge, there are only a limited number of models designed specifically for CRISPRi/a which generally provide all gRNAs that match the PAM sequence but do not account for efficiency scores^11^. Moreover, the few existing prediction models for CRISPRi/a have mainly been evaluated in promoter regions, and focus exclusively on transcriptomic impact ^10,12,13^.

Therefore, multiple critical gaps exist. First, putative enhancer regions are of primary interest in many studies but lack predictive models for epigenome editing experiments. Second, non-sequence features including comprehensive epigenetic annotations, the “essentiality” of nearest genes, and thermodynamic properties offer information complementary to sequence features. While these aspects have been somewhat explored, they have not yet been comprehensively incorporated into gRNA prediction for CRISPRi/a experiments. Finally, it is valuable to predict impact on multiple outcomes.

In this study, we present launch-dCas9, comprehensively predicting gRNA impact in multiple outcomes including cell fitness, wild-type abundance, gene expression, and generalizability across cell lines in CRISPRi screens. Methodologically, launch-dCas9 contains two types of prediction models, one employing a deep learning framework based on convolutional neural networks (CNN) and the other adopting the XGBoost^14^ method. We systematically benchmarked launch-dCas9 in multiple real CRISPRi datasets including large-scale genome-wide experiments with >1 million gRNAs designed for >100,000 candidate CREs, and evaluated its generalizability across multiple cell types.

## Results

### Launch-dCas9 Overview

Our launch-dCas9 (**Fig. 1**) constructs prediction models using features from multiple domains. Specifically, in addition to gRNA sequence information, we curated a list of functional annotations for each gRNA and its targeting DNase-I hypersensitive site (DHS) as input features. These annotations largely fall into three categories: thermodynamic properties, epigenetic marks and characteristics of nearest genes (**Methods**). For XGBoost, we manually generated mononucleotide and dinucleotide sequence features and combined them with functional annotations as model inputs. For CNN, we one-hot encoded the 20-bp protospacer sequences and employed four filters of varying sizes to capture information corresponding to different k-mers. The extracted sequence features were concatenated with functional annotations before passing through four fully connected layers (**Methods** and **Extended Data Fig. 1**).

**Fig. 1.**
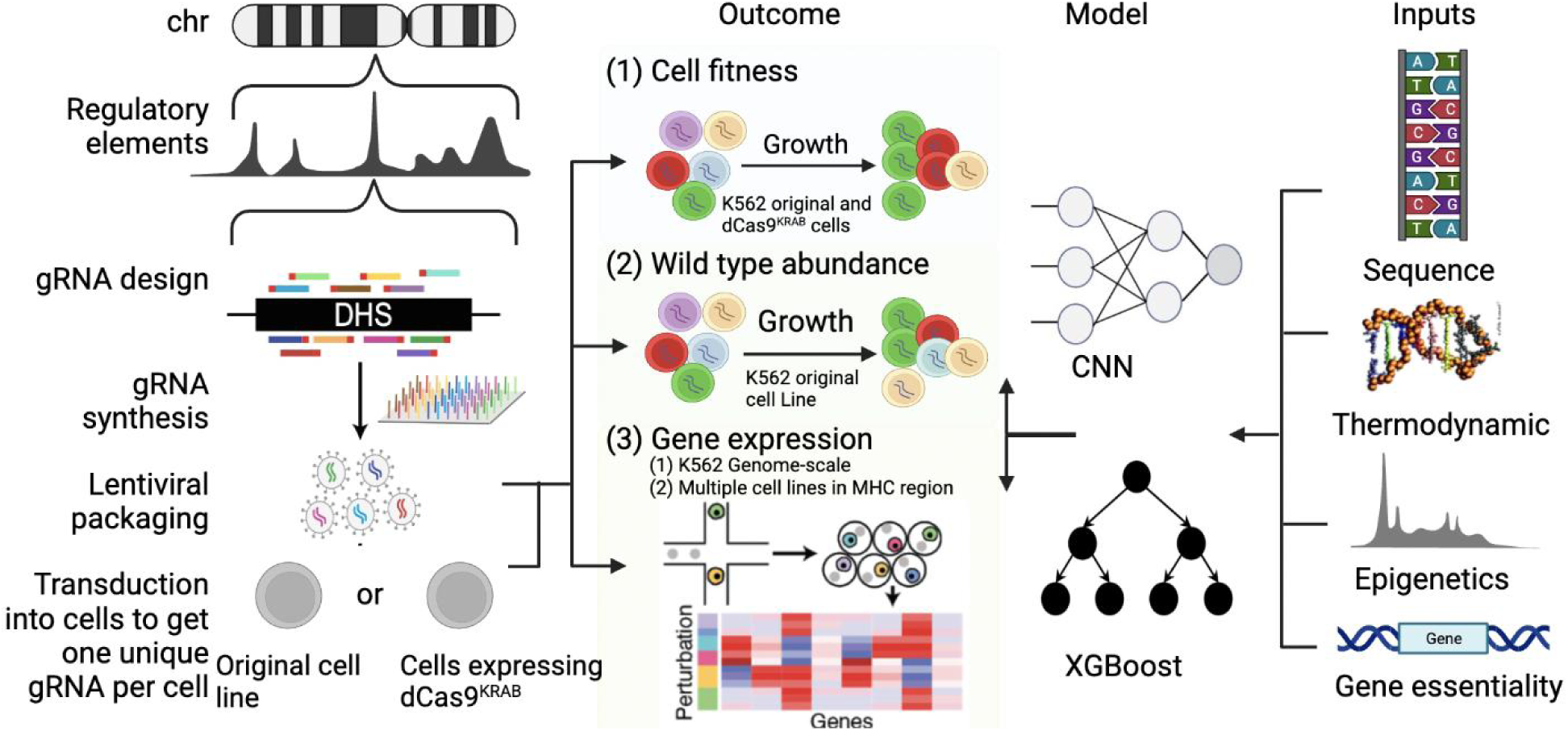
Overview of launch-dCas9. Adapted from (Klann et al., 2021). Our launch-dCas9 accepts four categories of input features: sequence-based features, thermodynamic quantifications, epigenetic annotations, and essentiality of nearest genes. It uses either CNN or XGBoost to predict outcomes including cell fitness, wild-type abundance, and gene expression changes. In real data, gene expression change is quantified using single-cell RNA-sequencing followed by CRISPRi perturbations. We used gene expression data at genome-scale in K562 cells and in the MHC region across multiple cell lines.

The number of DHSs and gRNAs considered for various types of analyses after preprocessing is shown in **Extended Data Fig. 2**. For training and evaluation of all models, we randomly split gRNAs into five folds according to their chromosome or DHS (**Supplementary Table 1**). Models were trained using four folds and tested on the held-out fold. Splitting by chromosomes avoids potential information leakage due to different gRNAs targeting the same DHS residing both in the training and test sets.

### Predicting Impact on Cell Fitness

We first trained our models using cell fitness data to predict the effects of gRNAs on cell growth. The whole genome CRISPR-dCas9-based epigenomic regulatory element screening (wgCERES) experiments we previously conducted involved >1 million gRNAs from >100,000 putative CREs, in human K562 leukemia cells^15^. We first categorized the gRNAs into significant or insignificant groups based on their FDR-adjusted p-values regarding their impact on cell growth (**Methods**). We assessed our launch-dCas9 models separately for gRNAs targeting promoter regions and those targeting enhancer regions.

### Promoter regions

In promoter regions, CNN and XGBoost achieved similar test-set AUC (mean AUC=0.800 for CNN and 0.803 for XGBoost) when all sequence and functional annotation features are used as model inputs (**Fig. 2a**). To characterize the importance of input features, we conducted ablation studies with different sets of model inputs. Models with only sequence information (mean AUC=0.707/0.711 for CNN/XGBoost) or only annotation features (mean AUC=0.770/0.776 for CNN/XGBoost) performed significantly worse than models utilizing all input features, suggesting that gRNA sequence and annotation features both provided important and complementary information for predicting the effect of gRNAs on cell growth.

**Fig. 2.**
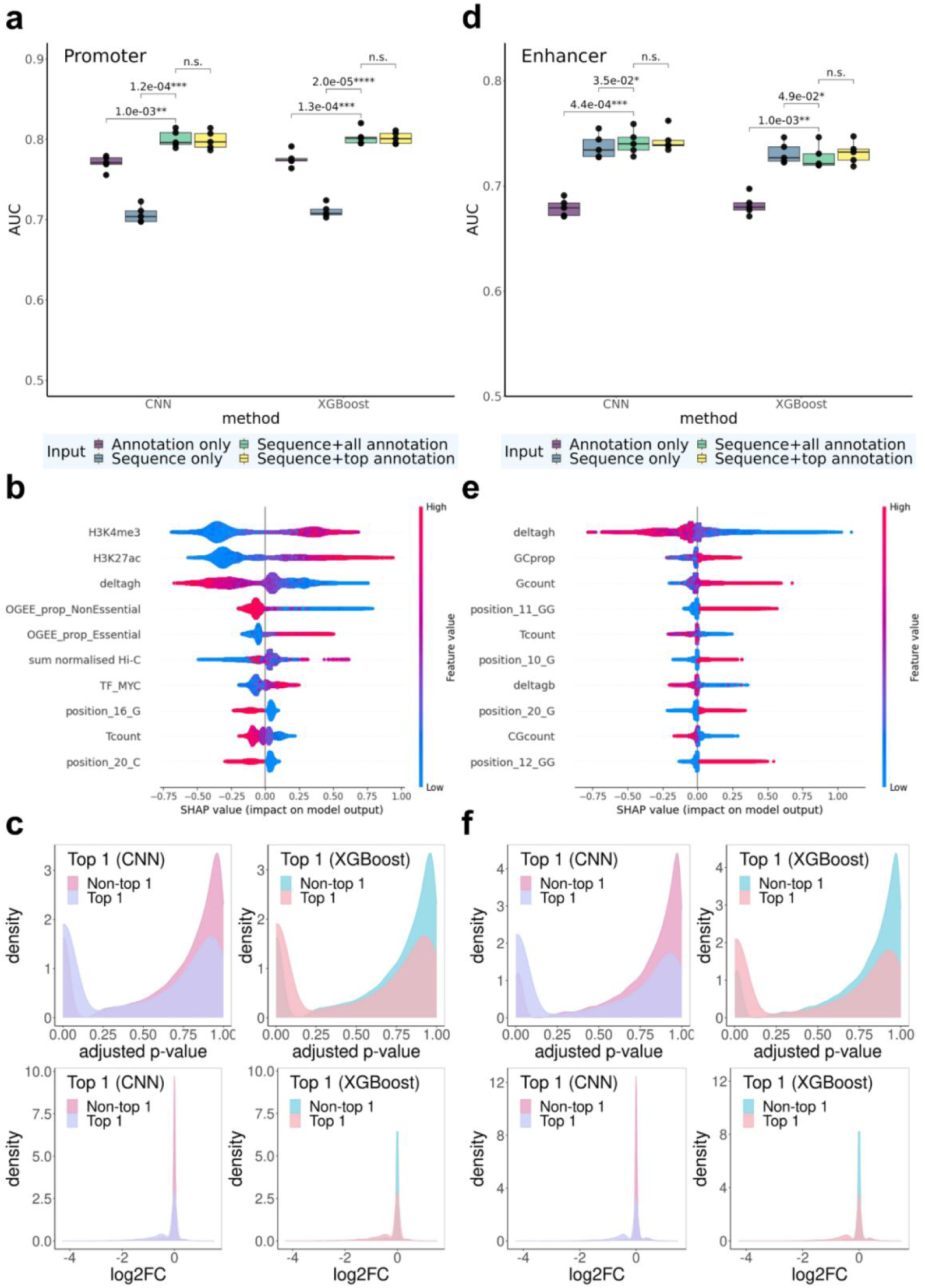
Predictive performance for cell fitness. **a-c,** gRNAs targeting promoter regions. **d-f,** gRNAs targeting enhancer regions. **a,d**: AUC for CNN and XGBoost models with different sets of input features. Statistically significant p-values from paired t-tests (across five folds) are listed above the bars. n.s.: not significant (p>0.05). **b,e**: SHAP summary plot that displays the top 10 most important features for XGBoost. **c,f**: Density plots of FDR-adjusted p-values and log_2_FC between the top 1 gRNA selected by CNN and XGBoost and the remaining gRNAs.

We further evaluated feature importance of the XGBoost model using SHapley Additive exPlanations (SHAP)^16,17^. SHAP values were used to quantify and visualize the contribution of each feature to model predictions. Similar to our findings from the ablation studies shown above, functional annotation features were critical, with the top six most important features all being functional annotations. Top features most predictive of gRNAs’ impact on fitness spanned all three categories of functional annotations in all five folds of our 5-fold cross validation assessments. Specifically, gRNAs with higher H3K27ac and H3K4me3 signals, lower *ΔG_H_* values, and higher essentiality of nearest genes^18^ were more likely to exert significant impact on cell fitness (**Fig. 2b**, **Supplementary Fig. 2a-d**). H3K4me3, an active promoter mark^19^, is expected to be associated with gRNAs’ impact on fitness in the direction observed. H3K27ac, albeit widely perceived as an active enhancer mark^20^, has been suggested to induce H3K4me3 enrichment around TSS and activate transcription when introduced to promoter regions^21^. The importance of H3K27ac in our models may also suggest that some promoters acting simultaneously as enhancers for other genes are likely to affect cell growth when perturbed. *ΔG_H_* measures gRNA-DNA hybridization free energy, and lower values indicate more efficient binding of a gRNA with its target DNA sequence and therefore higher probability of impactful perturbations. Gene essentiality (OGEE_prop_essential) was calculated as the proportion of cell lines in which the nearest gene was essential, therefore gRNAs targeting such regions may be more likely to affect cell fitness.

In view of the feature importance rankings, we additionally trained models utilizing sequence information and the top six functional annotation features (based on the SHAP values, see **Methods** for details) as inputs, and observed that they had similar performance compared with models utilizing all input features (mean AUC=0.799/0.805 for CNN/XGBoost compared to mean AUC = 0.800/0.803, p > 0.05), suggesting that those top annotational features captured almost all the annotation information essential for cell fitness prediction (**Fig. 2a**).

We conducted additional analyses to further assess the practical utility of launch-dCas9 in selecting gRNAs for future experiments. Specifically, for each DHS that has multiple gRNA choices (i.e., with ≥1 gRNAs significantly affecting cell fitness), we compared the predicted top 1 gRNA to the other gRNAs in the same DHS. Top 1 gRNAs predicted by both models (with the highest predicted probability of having effects) within each DHS had significantly lower experimentally derived FDR-adjusted p-values (adj. p) compared with other gRNAs (FDR <2.2e-16, Wilcoxon test). In particular, 36.4% and 36.6% of the top 1 predicted gRNAs had adj. p< 0.05 in our experiments for CNN and XGBoost, respectively; while only 14.5% of the other gRNAs had adj.p< 5% for CNN and XGBoost. Additionally, those top 1 gRNAs from both models had larger effect sizes on cell fitness (absolute values of log2-fold-change (log2FC)) compared with other gRNAs (p<2.2e-16, Wilcoxon test, **Fig. 2c**). The third quantile of |log2FC| from top 1 gRNAs was 0.519 for XGBoost and 0.510 for CNN, while that value from other gRNAs was only 0.074 for both models. We observed similar patterns for top 2 gRNAs (**Supplementary Fig. 4a**). We also compared the top observed gRNAs vs the others in their predicted probabilities of significance (**Supplementary Fig. 4c**) and observed similar patterns.

### Enhancer regions

For gRNAs targeting enhancer regions, CNN and XGBoost again achieved similar performance (mean AUC=0.742/0.730 for CNN/XGBoost) with all sequence and functional annotations as inputs (**Fig. 2d**). Ablation studies showed that annotation-only models performed significantly worse than others (mean AUC=0.679/0.682 for CNN/XGBoost), while sequence-only models achieved similar performances as the full model (mean AUC=0.738/0.731 for CNN/XGBoost). SHAP summary plots (**Fig. 2e**, **Supplementary Fig. 3**) reveal that top ranked features are all sequence-related features except the gRNA-DNA hybridization free energy ΔG_H_ and overall gRNA-DNA binding energy ΔG_B_. These results suggest that gRNA sequence information was more important for predicting effects in enhancer regions, and information captured by functional annotations are mostly contributed by chemical thermodynamic properties.

Again we observed no statistically significant differences between using all functional annotations and only top six selected annotations (mean AUC = 0.741/0.737 for CNN/XGBoost). Therefore, for all following analyses, we included only the top six annotations so that the trained models can be more easily applied in future experiments without the need of a huge number of annotations.

Analyses of top predicted gRNAs within each DHS showed consistent patterns as for promoter regions. The top 1 and top 2 gRNAs predicted by both models had significantly lower adj. p compared with other gRNAs (p<2.2e-16, Wilcoxon test). Specifically, 38.9% and 36.5% of the top 1 predicted gRNAs had adj. p< 5% in our experiments for CNN and XGBoost, respectively; while, for the remaining gRNAs, only 8.4% and 8.7% had adj. p< 5% for CNN and XGBoost, respectively. Additionally, the top 1 and top 2 predicted gRNAs exhibited larger |log_2_FC| than other gRNAs (p<2.2e-16, Wilcoxon test, **Fig. 2f**). The third quantile of |log_2_FC| from top 1 gRNAs was 0.46 for XGBoost and 0.47 for CNN, while that value from other gRNAs was only 0.05 for both models. Similar patterns were observed for the top 2 gRNAs as well (**Supplementary Fig. 4b**). These results demonstrate that launch-dCas9 indeed can not only predict DHSs that are likely to be effectively targeted and have effects on cell growth, but also select the most promising gRNAs within each DHS.

### Predicting gRNA Abundance in Wild-type Cells

The capacity to detect any significant gRNA effects depends on the number of sequencing-based counts of gRNAs identified in the baseline control group, i.e., among the wild-type cells. Certain gRNAs have lower packaging/transduction efficiency, or have intrinsic toxicity on cells when expressed on their own, and therefore show lower counts in both control and experimental groups due to the lack of cells carrying them, leading to insufficient statistical power. Therefore we utilized wild-type cells (cells without dCas9-KRAB introduced) to investigate whether launch-dCas9 can accurately predict which gRNAs are more likely to produce low counts. Since epigenomic features of target sites are not expected to affect gRNA packaging or expression efficiency and our gRNAs mostly target enhancer regions (**Extended Data Fig. 2**), we only used sequence information as model inputs. Both models achieved a strong correlation (mean Spearman correlation across fivefolds: 0.799/0.689 for CNN/XGBoost) between predicted and true counts (**Fig. 3a-b**). The worse performance of XGBoost is potentially due to the fact that manually curated inputs to XGBoost only contain mononucleotide and dinucleotide sequence information, missing features accounting for longer k-mers, while CNN captures longer k-mer information by including kernels of sizes 3, 5 and 7 nucleotides (nt). To test this hypothesis, we conducted ablation experiments removing different kernels from the CNN architecture (**Fig. 3a** and **Supplementary Fig. 5**). CNN without kernels of sizes 5 and 7 performs significantly worse than the original CNN, and the performance further decreases to similar levels as XGBoost when kernels of size 3 are eliminated as well. This further demonstrates that complex features accounting for longer k-mer information can be important in predicting wild-type counts.

**Fig. 3.**
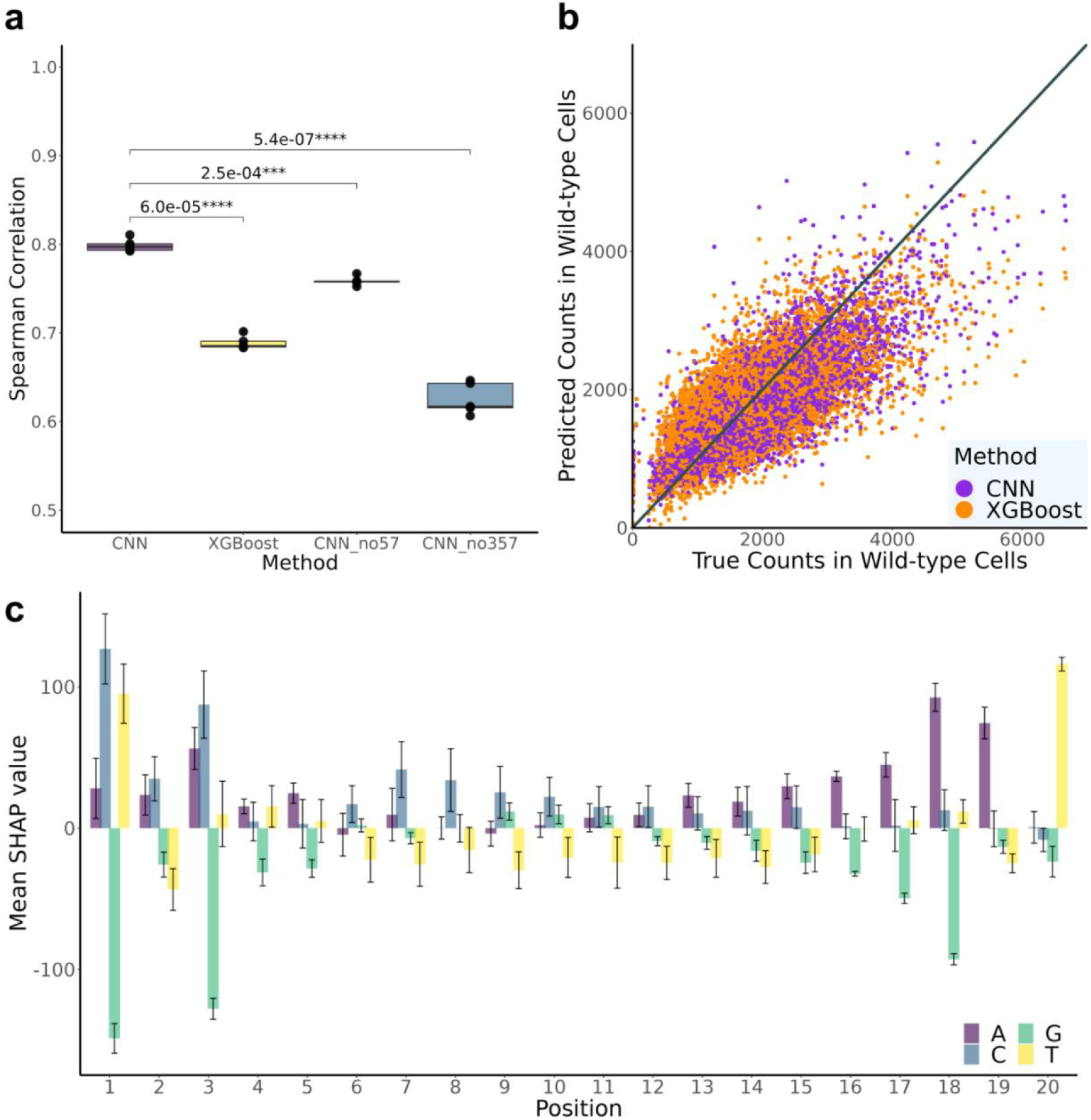
Prediction performance for counts in wild-type cells. **a,** AUC for XGBoost and CNN models with different convolutional layer structures. CNN_no57: CNN without kernels of size 5 and 7 nt; CNN_no357: CNN without kernels of size 3, 5, and 7 nt. P-values from paired t-tests are listed above the bars. **b,** Scatterplot of predicted values versus true counts in wild-type cells. **c,** Mean SHAP values for different input sequence positions. Error bars represent ±one standard deviation across five folds.

We further utilized DeepSHAP^16^ to pinpoint which positions of the input sequence are most important for determining counts in wild-type cells (**Fig. 3c** and **Supplementary Fig. 6**). Nucleotides at the first and last three positions of the gRNA sequence influence model predictions the most. Guanine (G) at position 1, 3, and 18 all greatly contribute to lower counts; while cytosine (C) at position 1 and 3, thymine (T) at position 1 and 20, and adenine (A) at position 18 and 19 all contribute to higher counts. These findings provide important insights for future gRNA designs in CRISPRi experiments.

### Predicting impact on nearby gene expression levels from single-cell CRISPRi experiments

We have previously performed single cell CERES (scCERES) experiments where we perturbed 3,029 DHS with one gRNA per DHS^15^. Differential gene expression results were obtained on genes ±1Mb of each DHS (**Methods**). To predict gRNAs’ impact on gene expression, we trained and assessed launch-dCas9 prediction models to predict whether a gRNA affects expression of any genes in the ±1Mb neighborhood. For this purpose, we selected the most significant gRNA-gene pair for each gRNA. Similar to the modeling framework for predicting cell fitness, we categorized gRNAs as having significant effects or not on the expression of any nearby gene (**Methods**) and utilized sequence information and top functional features to train prediction models. CNN and XGBoost achieved similar AUC (mean AUC = 0.630/0.635 for CNN/XGBoost) when all sequence and top functional annotation features are used (**Fig. 4a**). Ablation studies with different sets of model inputs show that models with only sequence information (mean AUC = 0.518/0.513 for CNN/XGBoost) performed significantly worse than those utilizing all input features, while annotation-only models achieved similar or even better performance as the full model (mean AUC = 0.602/0.660 for CNN/XGBoost). Among all the annotation features, the most important features are H3K27ac, ATAC-seq, and H3K4me3 (**Supplementary Fig. 6**). Therefore, we conclude that gRNA annotation features are adequate to provide important information for predicting the effect of gRNAs on nearby gene expression.

**Fig. 4.**
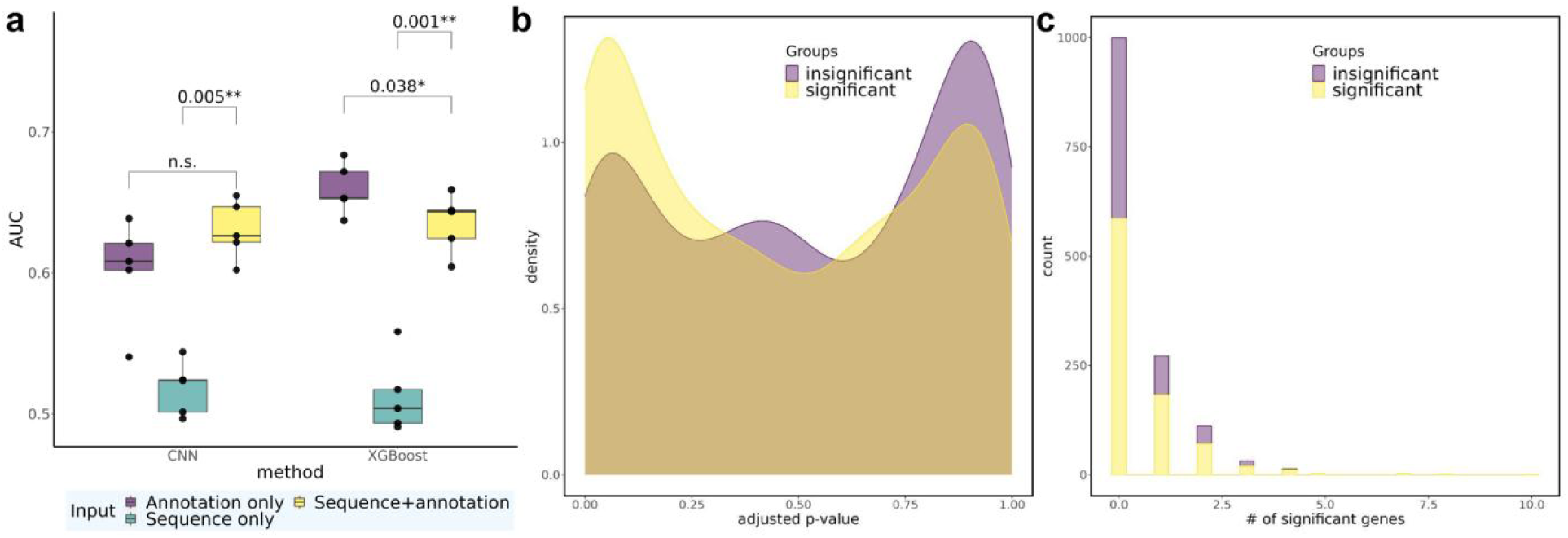
Predictive performance on gene expression and relationship between transcriptomic impact and cell growth. **a,** AUC for CNN and XGBoost models with different sets of input features in scCERES. Statistically significant p-values from paired t-tests are listed above the bars. n.s.: not significant (i.e., p > 0.05). **b,** Density plot illustrating the distribution of FDR-adjusted p-values for the two groups of gRNAs, based on their observed impact (significant or insignificant) on cell fitness. **c,** Histogram illustrating the distribution of the number of genes significantly influenced by a gRNA, separately for the two groups of gRNAs, again based on their impact on cell fitness.

### Relationship between gene expression and cell fitness

To investigate gene regulation mechanisms, we examined whether gRNAs in wgCERES exhibit any enrichment in scCERES. We again divided gRNAs into two groups based on whether they had significant impact on cell fitness and found that gRNAs belonging to the significant group had a higher density of small FDR-adjusted p-values in scCERES experiments. For example, among gRNAs having significant impact on cell fitness, 23.4% had significant effect on gene expression (adj. p<0.2), while among gRNAs having no significant impact on fitness, 17.1% had significant effect on gene expression (adj. p< 0.2; **Fig. 4b**). The difference between the two groups is likely not large, both because all the sgRNAs in this library have been previously shown to significantly impact cell fitness and we chose the most significant gene for each sgRNA. Furthermore, we discovered a significant association between the significance of a gRNA’s impact on cell fitness and its impact on expression of genes in the neighborhood (**Fig. 4c**, odds ratio = 1.44, Chi-square test *p* = 0.003, detailed in **Methods** and **Supplementary Table 2a**). Additionally, there exists a significant enrichment between significant impact on cell fitness and influence on expression of >3 genes (odds ratio =6.77, Chi-square test *p* = 0.006, **Supplementary Table 2b**). This suggests a strong positive association between impact on cell fitness and on gene expression, particularly through regulation of multiple genes in the neighborhood.

### Guide RNA effect prediction models are reasonably generalizable in the MHC region across cell lines

To assess the generalizability of our prediction models across different cell lines, we used scCERES datasets designed to target the same 581 regions. The majority of these regions are open chromatin areas in at least one cell type, while the others are regulatory elements previously identified in a bulk CRISPR screen. These regions were targeted across K562, induced pluripotent stem cells (iPSCs), and human iPSC-derived neural progenitor cells (NPCs) within the Major Histocompatibility Complex (MHC) region (**Methods, Extended Data Fig. 2**). Here, data were split into 5 folds according to target regions. Additionally, we applied models trained from the whole genome scCERES in K562 to evaluate its transferability to the MHC region. Our findings indicate that the training models have the best performance when applied to the same cell line (**Fig. 5a**). Interestingly, the most important factor that influences performance is the test cell line itself, specifically K562 and iPSC are overall relatively easier to predict (median AUC = 0.71 for K562, median AUC = 0.67 for iPSC) than NPC (median AUC = 0.61). Given any particular test cell line, the relative loss when using models trained from unmatched cell lines is not large, particularly for CNN where there is no significant difference among all trained models for any test cell line. Overall, our results suggest that gRNA effect prediction models are reasonably generalizable in the MHC region across the three cell lines assessed.

**Fig. 5.**
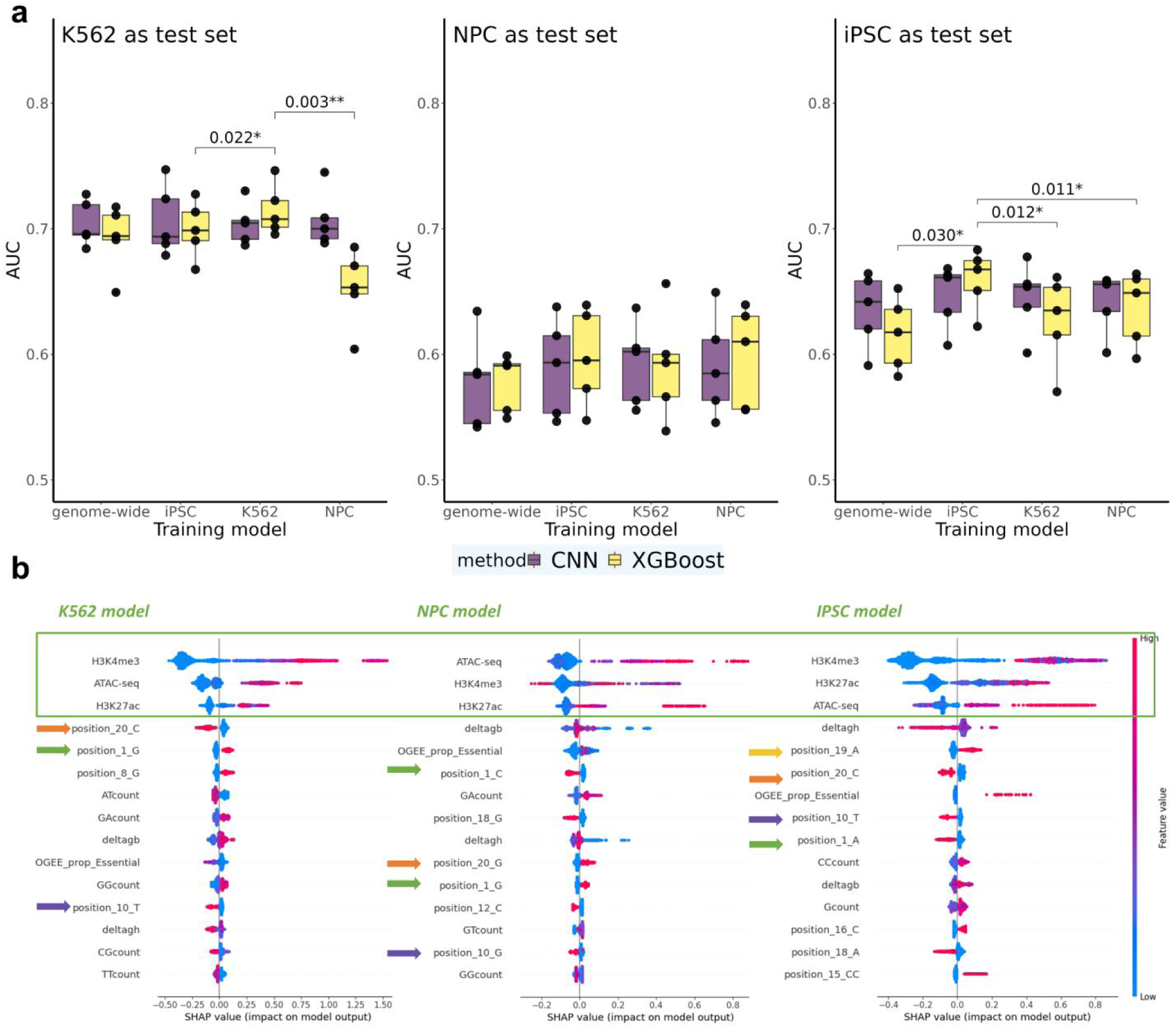
Predictive performance for nearby gene expression in the MHC region across different cell types. **a,** AUC for CNN and XGBoost models across multiple cell lines in the MHC region. Statistically significant p-values from paired t-tests are listed above the bars. Tests were performed comparing models trained on the same cell line as test set versus models trained on other cell lines. Non-significant comparisons (p>0.05) are omitted. **b,** SHAP summary plot for XGBoost models, highlighting the top 15 most significant features in each cell line model.

SHAP plots from XGBoost models across the three cell lines may explain the reason for generalizability (**Fig. 5B**). These plots consistently revealed the same top three most important features across all three cell lines. Additionally, among sequence-derived features, bases at positions 1, 10, 19, and 20 all rank among the top 10. The similarity in feature importance across cell lines is consistent with our observed model transferability. In addition, different mononucleotides and dinucleotides at those top positions may explain the lower generalizability of XGBoost compared to CNNs, e.g., position_1_G vs position_1_C vs position_1_A. This is because CNNs are better equipped to handle position-correlated information, while XGBoost requires pre-engineered features for the same positions.

## Discussion

In this study, we present launch-dCas9, which employs CNN and XGBoost frameworks, to comprehensively predict gRNA impact on cell fitness, wild-type abundance, and single-cell gene expression in CRISPRi experiments, and to systematically assess the contributions of sequence and functional annotation features to the prediction models. Our launch-dCas9 will facilitate easy and efficient design and selection of top gRNA protospacers within each DHS, as well as across the genome, in multiple ways, including revealing the most important features specific to different tasks, and selecting gRNAs with higher wild-type abundance that allows for more powerful statistical testing of gRNA impact. Our launch-dCas9 models show reasonable transferability across three cell lines evaluated, suggesting that models trained on other cell lines or genome-wide screens can serve as an alternative way to help optimize CRISPR on-target design in cell types where we have no or limited CRISPRi experiments to train reliable models.

In this study, we discovered that the sequence position preferences in CRISPRi differ from those in CRISPR/Cas9 experiments, consistent with findings reported in Xu et al^10^. For instance, while nucleotides proximal to the PAM are important for both CRISPR/Cas9 and CRISPRi/a screens^23,24^; positions 1 and 3 are particularly predictive only for CRISPRi screens. Also, the efficiency of gRNA in CRISPRi appears to rely less on the sequence context alone. Such differences can be attributed to the fact that CRISPRi utilizes a unique mechanism to disrupt gene function without causing DNA cleavage.

Cell growth can be affected through multiple potential mechanisms, among which a crucial one is through perturbations in gene expression levels. In this study we have briefly explored the relationship between gRNAs impacting cell fitness and gene expression, and discovered that gRNAs impacting cell growth are more likely to lead to changes in gene expression. However we are underpowered to identify the causal genes or the cascade of expression changes in certain orchestrated pathways. Future comprehensive experiments may shed more light on the regulatory element-gene pairs that work together to affect cell fitness.

Our study interestingly revealed that input features rank differently for different tasks. For instance, functional annotation features that substantially affect performance when predicting impact on expression levels of nearby genes, are largely dispensable when predicting impact of gRNAs on cell fitness. Our results provide practical guidelines regarding which features to prioritize according to the task of interest.

Our launch-dCas9 leverages XGBoost and CNN to construct prediction models. We comprehensively compare the performance of the two methods for different tasks. Generally, their performance is similar which is consistent with previous findings^24^. However, XGBoost tends to outperform in tasks where functional annotations are more important than sequence information, whereas CNN excels in capturing longer sequence information. When it comes to transferability across cell types, CNN appears to have relatively better generalizability than XGBoost. This is especially true for cell types that are more challenging to train (e.g., NPC).

The large number of gRNAs in our experiments, particularly the >1 million gRNAs across >100,000 DHSs from wgCERES, renders it possible to train deep learning models. However, for many experiments, due to various reasons including but not limited to small number of gRNAs, insufficient number of single cells, or low wild type expression levels, the number of labeled outcomes (e.g., gRNAs with high-confident significant or insignificant effect on a certain outcome) can be rather small. Rather, most gRNAs fall into the gray zone where we have insufficient evidence to conclude as either significant or insignificant with high confidence.

Therefore, semi-supervised models^25^ that accommodate a large number of unlabelled data (in the gray zone) warrant future explorations beyond the scope of this manuscript.

In conclusion, we present launch-dCas9, as the first prototype model for comprehensively optimized gRNA design in CRISPRi experiments that we believe will be broadly valuable for many investigators.

## Material and methods

### Convolutional Neural Network structure

We developed a CNN model based on two previous models developed for CRISPR-Cas9 gRNA efficiency predictions^4,6^. The input 20-bp protospacer sequences were first converted into four-by-twenty binary matrices through one-hot encoding. Launch-dCas9-CNN starts with a one-dimensional convolutional layer consisting of four different kernel sizes with varying numbers of filters similar to an inception module: 50, 100, 70 and 40 filters for kernels of 2, 3, 5, and 7 nt respectively. All components are followed by a max pooling layer of size 2 and a dropout layer with dropout rate of 0.4, with the exception of kernels of 2 nt which uses a stride of 2 and no pooling layers. The outputs from the four components are flattened and concatenated before passing through a fully connected layer with 80 nodes and a dropout rate of 0.3. The 80-dimensional outputs represent extracted sequence features. We additionally applied a size-1 kernel to the input one-hot encoded sequences to extract 20-dimensional mononucleotide information. These two are concatenated with the vector of functional annotation inputs before passing through three fully connected layers with 80, 60, 40 nodes respectively and a dropout rate of 0.4. A final fully connected layer with one output node follows. For all layers except the last one, we use Rectified Linear Unit (ReLU) as activation functions. When training CNN models, we used Adam optimizer with a learning rate of 0.0001 for cell fitness and wild-type counts, and a learning rate of 5e-5 for gene expression predictions. Missing data were preimputed with zero imputation except variable ‘ploidyZhou’ for which we used mode imputation. All binary models (for cell fitness and gene expression tasks) were trained using binary cross entropy and all continuous models (for wild-type counts) were trained by minimizing mean square error (MSE). We used a batch size of 256 for cell fitness predictions and 512 for the other two tasks.

### eXtreme Gradient Boosting(XGBoost) for features analysis

Due to XGBoost’s inability to handle sequence information directly as CNN, we utilized a bag-of-words (BoW) model to tokenize each gRNA and represent it as an input vector. Our first step involves designing a vocabulary based on the position and bases of the gRNA sequence. Each gRNA has a 20-base pair protospacer, resulting in a total of 80 (20 * 4) unique features based on the combination of position and mononucleotide and 304 (19 * 4 * 4) unique features based on the combination of position and dinucleotide. We then consider the frequency of each mono- and dinucleotide within a gRNA, resulting in 20 (4+4*4) position-independent unique features.

Consequently, aside from functional annotations, we include 404 sequence features in our XGBoost analysis. We used 5-fold cross-validation to select the best model from 57 different configurations of regularization parameters and hyperparameters within the training data. The hyperparameters we searched over included the maximum depth of a tree (chosen from [3, 5, 7, expanded until all leaves are pure]), the minimum sum of instance weight (hessian) needed in a child (chosen from [8, 12, 16, 20]), the subsample ratio of the training instances (chosen from [0.5, 0.6, 0.7, 0.8, 0.9, 1]), the subsample ratio of columns when constructing each tree (chosen from [0.5, 0.6, 0.7, 0.8, 0.9, 1]), the L1 regularization term on weights (chosen from [10, 30, 50, 70, 90]), and the step size shrinkage (chosen from [0.3, 0.2, 0.1, 0.05]). To select the best model, we evaluated the performance (using AUC) on the whole genome training dataset for cell fitness and gene expression tasks, and root mean square error for the wild-type counts task. After selecting the best model, we applied its optimal hyperparameters to each fold, which were split by chromosome, to train the model and generate final predictions on the test sets for each fold.

### Datasets to derive functional annotation features

The aligned sequence read files (in BAM format) of all functional annotation marks (H3K4me3, ATAC-seq, H3K27ac, CHIP-seq of transcription factors) are downloaded from the ENCODE Portal (http://encodeproject.org) unless stated otherwise. The ENCODE accession numbers are listed in Supplementary Table 3. We used the human genome assembly hg19 for all analyses.

The NPC ATAC-seq data were generated by Inoue et al. (2019)^26^. We downloaded the FASTQ files provided by the authors and processed them using the ENCODE pipeline to obtain the BAM files. We only used files corresponding to the last time point (72hr). All the defination of annotation features are listed in **Supplementary Text**.

### K562

H3K4me3 ChIP-seq data were produced by the Stamatoyannopoulos lab for human P5 wild type. ATAC-seq data was produced by the Snyder lab. H3K27ac ChIP-seq data was produced by the Bernstein lab. We also downloaded the ChIP-seq data for the following transcription factors (TF): GATA2, TAL1, MYC, all from the Myers lab.

### WTC11 iPSC

ATAC-seq data was produced by the Shen lab while H3K4me3 and H3K27ac histone ChIP-seq data were produced by the Bernstein lab.

### NPC

H3K4me3 and H3K27ac CHIP-seq data were produced by the Ren lab.

### Counting mapped reads for functional features

Firstly, bam files were indexed by Rsamtools^27^. Rsubread^28^ then took mapped reads as input and assigned them to the tiled genomic features with 1kb width on hg38. Positions from all reads, excluding those on chromosome Y, were lifted from the hg38 to the hg19 assembly.

### Data processing and label definition

For cell fitness outcome in wgCERES, gRNA counts were extracted in both dCas9^KRAB^ and control (no dCas9^KRAB^) conditions. Differential expression analysis was performed using DESeq2^32^. The gRNA abundance outcome in the wgCERES assay was calculated as the average gRNA counts based on amplicon sequencing across four replicates in wild-type cells from screens in K562. For gene expression outcome in scCERES and MHC experiments, logFC of transcript counts between cells containing the gRNA versus all other cells were derived and MAST^33^ was used to identify that significant differences. Among all the experiments, gRNAs with FDR-adjusted p-values smaller than 0.05 are labeled as having significant effects on the corresponding outcome (be it cell fitness, wild-type counts, or gene expression of nearby genes), while gRNAs with FDR-adjusted p-values larger than 0.2 are labeled as not having significant effects. To avoid potential mislabeling of borderline significant gRNAs, we did not use gRNAs with a FDR-adjusted p-value between 0.05 and 0.2 in either model training or testing. To focus only on gRNAs with reasonable power of detecting significant effects for the cell fitness outcome, we plotted the total counts versus -log10(*adj.p-value*) for each gRNA and decided to remove all gRNAs with total counts < 125. The promoter regions for the cell fitness outcome were determined using the annotatePeak function from ChipSeeker^34^, while all other regions were categorized as enhancers for the wgCERES screen. As CNNs are sensitive to the range of input features, for all the continuous features, we performed winsorization by capping extreme values at the 95th percentile to reduce the impact of outliers. Then, we fed the winsorized features to both CNN and XGBoost.

When evaluating the effectiveness of launch-dCas9 in choosing gRNAs for future studies, we focused on DHS regions that had at least three significant gRNAs. We defined the top 1 gRNA as the one having the highest predicted probability among gRNAs in the same DHS and similarly for the top 2 gRNAs.

### Comparing between different experimental results

P-values were calculated using paired t tests from the function ‘t_test’ in R package ‘rstatix’ and FDR adjusted using the Benjamini-Hochberg^35,36^ method.

### Association between gene expression and cell fitness

We conducted two analyses to determine whether the effect of gRNA on cell fitness is associated with gene expression. We identified all the gRNAs that were present in both wgCERES and scCERES screens. As the scCERES screen was based on the results of wgCERES, the significant group contained more gRNAs than the insignificant group. We had 881 gRNAs from the significant group and 557 gRNAs from the insignificant group. Firstly, we only included the most significant FDR-adjusted p-value of gene expression for analysis and plotted the density difference of adjusted p-value between the two groups of gRNAs, split based on their impact on cell fitness (**Fig. 4b**). Secondly, we included all gRNA-gene pairs and performed a chi-square test to quantify the association between the impact of gRNA on cell fitness and its effect on the expression of one or more gene(s). We identified gRNAs that had a significant impact on gene expression with *adj.p-value*<0.2. A table was constructed by grouping the gRNAs based on whether they had a significant impact on cell fitness and the number of significant genes (**Fig. 4c, Supplementary Table 2B**).

## Author contributions

W.M and T.L. developed the method and performed the computational experiment. L.B, T.K, M.W performed the data expermentation. L.B., T.K., and M.W. conducted the data experimentation. A.B. and J.B. were responsible for cleaning and preprocessing the raw data. G.C., P.S., C.G., M.L., and Y.L. offered crucial guidance and oversight throughout the study. The manuscript was primarily authored by W.M., T.L., and Y.L., with invaluable input and approval on the final draft from all contributors.

## Data and code availability

Launch-dCas9 is implemented in Python with the source code available at https://github.com/Wancen/launch-dCas9. All custom Python scripts to generate result figures for this paper are available on GitHub (at https://github.com/TianyouLuo/gRNA_prediction).

## Extended Data Figure

**Extended Data Fig. 1.**
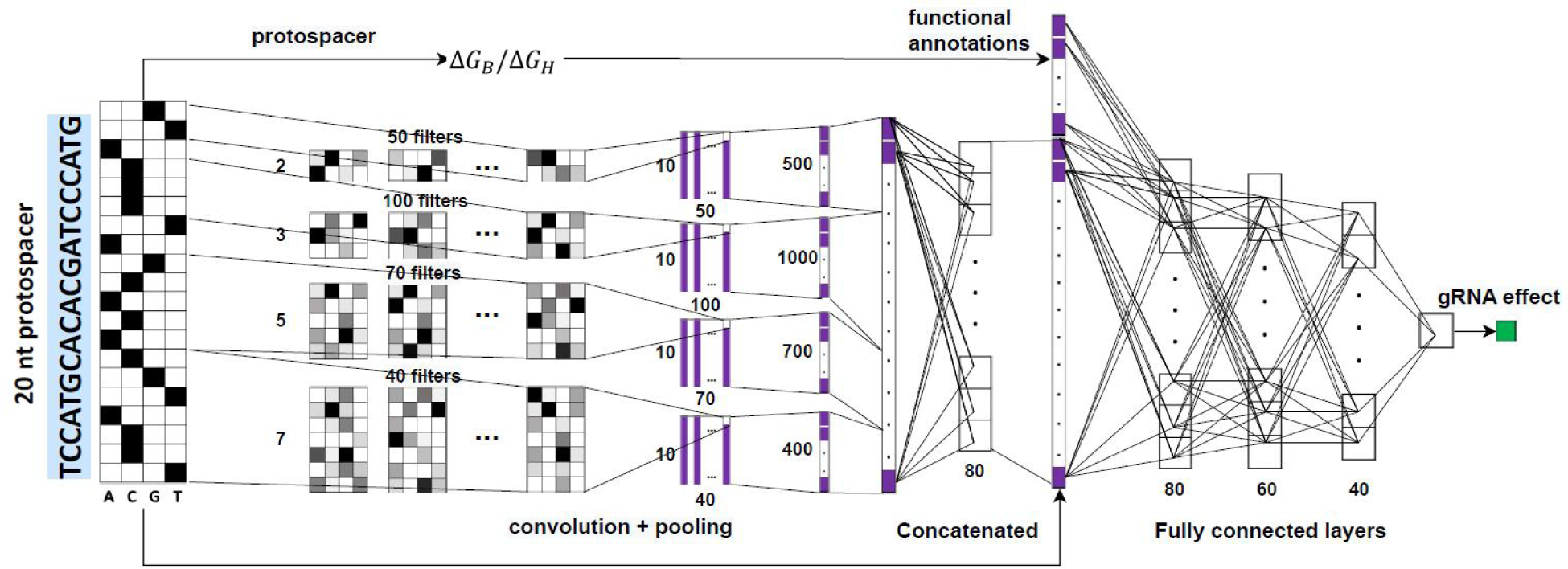
CNN architecture. The inputs to the CNN are the one-hot encoded 20mer, the thermodynamic related features(ΔG_B_, ΔG_H_), functional annotations, gene essentiality related features. Figure is created based on Fig. 2a from Xiang et al^6^

**Extended Data Fig. 2.**
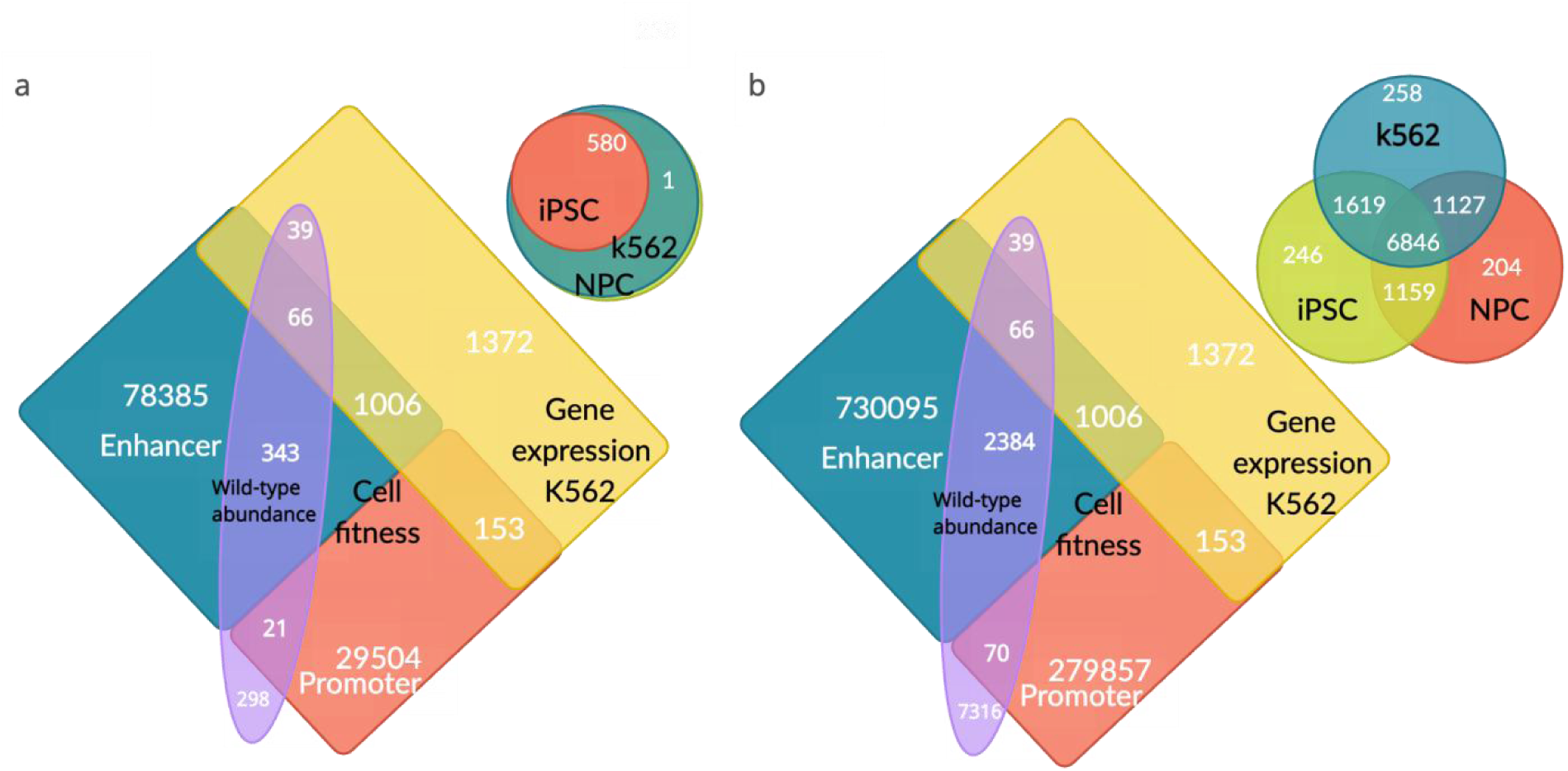
Venn Diagrams Illustrating Target Regions, DHSs, and gRNAs Employed in Various Prediction Models and Analyses. **a:** The bottom-left quadrant shows DHSs used in both wgCERES and scCERES screens, while the top-right quadrant displays target regions from the MHC experiments. **b:** The bottom-left quadrant features gRNAs employed in both wgCERES and scCERES screens, and the top-right quadrant presents gRNAs used in MHC experiments.

