## Supplementary Text, Figures, and Tables for "Machine learning methods for predicting guide RNA effects in CRISPR epigenome editing experiments"

### Supplementary Files

*Wancen Mu et al.*

#### Supplementary Text

**Description of annotation features:**

Our methodology has been fortified by the careful selection of top annotation features guided by their SHAP values derived from cell fitness prediction tasks in enhancer and promoter regions. From a comprehensive list of annotation features given below, we prioritized six annotations based on their prominence and relevance in the studied context. These annotations, namely H3K4me3, ATAC-seq, H3K27ac, Deltagh, Deltagb, and OGEE_prop_Essential, were chosen for their significant roles and contributions to the gene regulatory mechanisms and cellular functionalities.

**H3K4me3**: H3K4me3 CPM per 1kb for each DHS. If a DHS entirely overlaps two (or more) tiled genomic regions with 1kb width, the summation is shown.
**ATAC-seq**: ATAC-seq CPM per 1kb for each DHS. If a DHS entirely overlaps two (or more) tiled genomic regions with 1kb width, the summation is shown.
**H3K27ac:** H3K27ac CPM per 1kb for each DHS. If a DHS entirely overlaps two (or more) tiled genomic regions with 1kb width, the summation is shown.
**TF_GATA2**: GATA2 CPM per 1kb for each DHS. If a DHS entirely overlaps two (or more) tiled genomic regions with 1kb width, the summation is shown.
**TF_TAL1**: TAL1 CPM per 1kb for each DHS If a DHS entirely overlaps two (or more) tiled genomic regions with 1kb width, the summation is shown.
**TF_MYC**: MYC CPM per 1kb for each DHS. If a DHS entirely overlaps two (or more) tiled genomic regions with 1kb width, the summation is shown.
**Deltagb:** The free energy metric that quantifies binding affinity of the Cas9–gRNA–DNA complex according to Alkan et al. 2018[29]
**Deltagh:** The free energy contribution of gRNA–DNA hybridization according to Alkan et al. 2018[29]
**olap_number:** The number of overlapping Hi-C (High-throughput chromosome conformation capture) pairs[30] with the 20 bp gRNA target site for each gRNA
**sum_log10fdr:** The total log10-transformed FDR-corrected p-value of the overlapping Hi-C pairs with the 20 bp gRNA target site for each gRNA

**prom_olap_number:** The number of overlapping Hi-C pairs, where one side is located in the promoter region and the other side is within the 20 bp gRNA target site. The promoter regions of genes were obtained from the gencode.v28lift37.txt.

**prom_sum_log10fdr:** The total log10-transformed FDR-corrected p-value of the overlapping Hi-C pairs, where one side is located in the promoter region and the other side is within the 20 bp gRNA target site for each gRNA. The promoter regions of genes were obtained from the gencode.v28lift37.txt.

**ploidyZhou**: Large scale ploidy of the region according to Zhou et al. 2019 (NA value if ploidy of the region was not reported or if gRNA overlap two regions with different ploidy)

**LossHetZhou**: True if region lost heterozygosity according to Zhou et al. 2019, False otherwise

**SV_Zhou**: True if structural variant overlap gRNA according to Zhou et al. 2019, False otherwise

**SNV_Zhou**: True if single nucleotide variant overlap gRNA according to Zhou et al. 2019, False otherwise

**conserved_score**: select the max score if gRNA overlaps two or more conserved regions according to Zoonomia Project[31].

**Conserved**: True if gRNA overlap conserved regions (>top90% percentile of 10bp regions) according to di Iulio et al. Nat Gen 2018. False otherwise

**Gquad_n_overlap_same_strand**: Number of bases of the gRNA overlapping a G4 quad on the same strand (0 if no overlap between gRNA and G4 quad)

**Gquad_n_overlap_other_strand**: Number of bases of the gRNA overlapping a G4 quad on the other strand (0 if no overlap between gRNA and G4 quad)

**distance**: distance of the gRNA to the closest protein coding gene on the same strand (gene definition from gencode.v29lift37). Negative if before TSS, positive if after the TES, 0 if within gene boundaries.

**medianRNAseqTPM**: Median of the mean TPM of 4 ENCODE K562 RNA-seq experiments. The 4 experiments are "ENCSR000AEM", "ENCSR000AEO", "ENCSR000CPH", "ENCSR545DKY"), each with 2 technical replicates. We first took the mean TPM across replicates for each experiment, then took the median of the mean TPMs across the 4 experiments for genes measured in all 4)

probIntolerantLoF: probability that the gene is intolerant to loss of function mutation (from: Lek et al. Nature, 2016)

**numTKOHits_Hart**: number of cell lines in which the gene is essential (from Hart et al., Cell 2018)

**HartEssential**: True if genes is essential in more than 2 of the Hart et al. cell lines (their definition of essential genes, genes with 1 or 2 could be defined as conditionally essential)
**OGEE_n_Essential**: number of cell lines in which the gene is essential according to the OGEE database (http://ogee.medgenius.info)
**OGEE_n_NonEssential**: number of cell lines in which the gene is non-essential according to the OGEE database (http://ogee.medgenius.info)
**OGEE_n**: number of cell lines in which the gene was tested for essentiality according to the OGEE database (http://ogee.medgenius.info)
**OGEE_prop_Essential**: proportion of cell lines in which the gene is essential according to the OGEE database (http://ogee.medgenius.info)
**OGEE_prop_NonEssential**: proportion of cell lines in which the gene is non-essential according to the OGEE database (http://ogee.medgenius.info)
**cancer_census_tier**: cancer tier (https://cancer.sanger.ac.uk/). Tier1: To be classified into Tier 1, a gene must possess a documented activity relevant to cancer, Tier2: genes with strong indications of a role in cancer but with less extensive available evidence. Value set to 0 for non-cancer genes.
**cancer_census_tissue_type**: L: leukemia/lymphoma, E: epithelial, M: mesenchymal, O: other.
**sum normalised Hi-C**: sum of normalised Hi-C for each DHS (from ABC file). Values are only shown for gRNA entirely within the DHS. If a gRNA entirely overlaps two extended DHS, the mean is shown.

#### Supplementary Figure

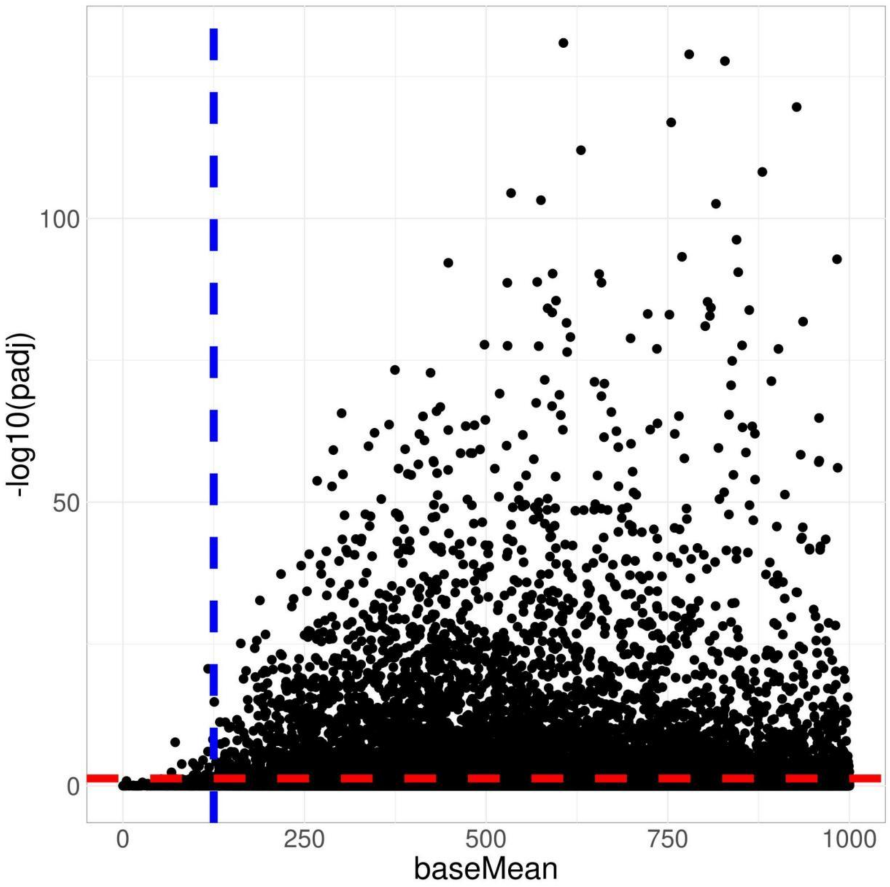


***Supplementary Fig. 1* | *Scatter plot of total counts versus -log10(adj.p-value) for all gRNA in wgCERES screen. The blue dashed line represents the threshold of total counts at 125, while the red dashed line represents the threshold of -log10(adj.p-value) at 0.05.***

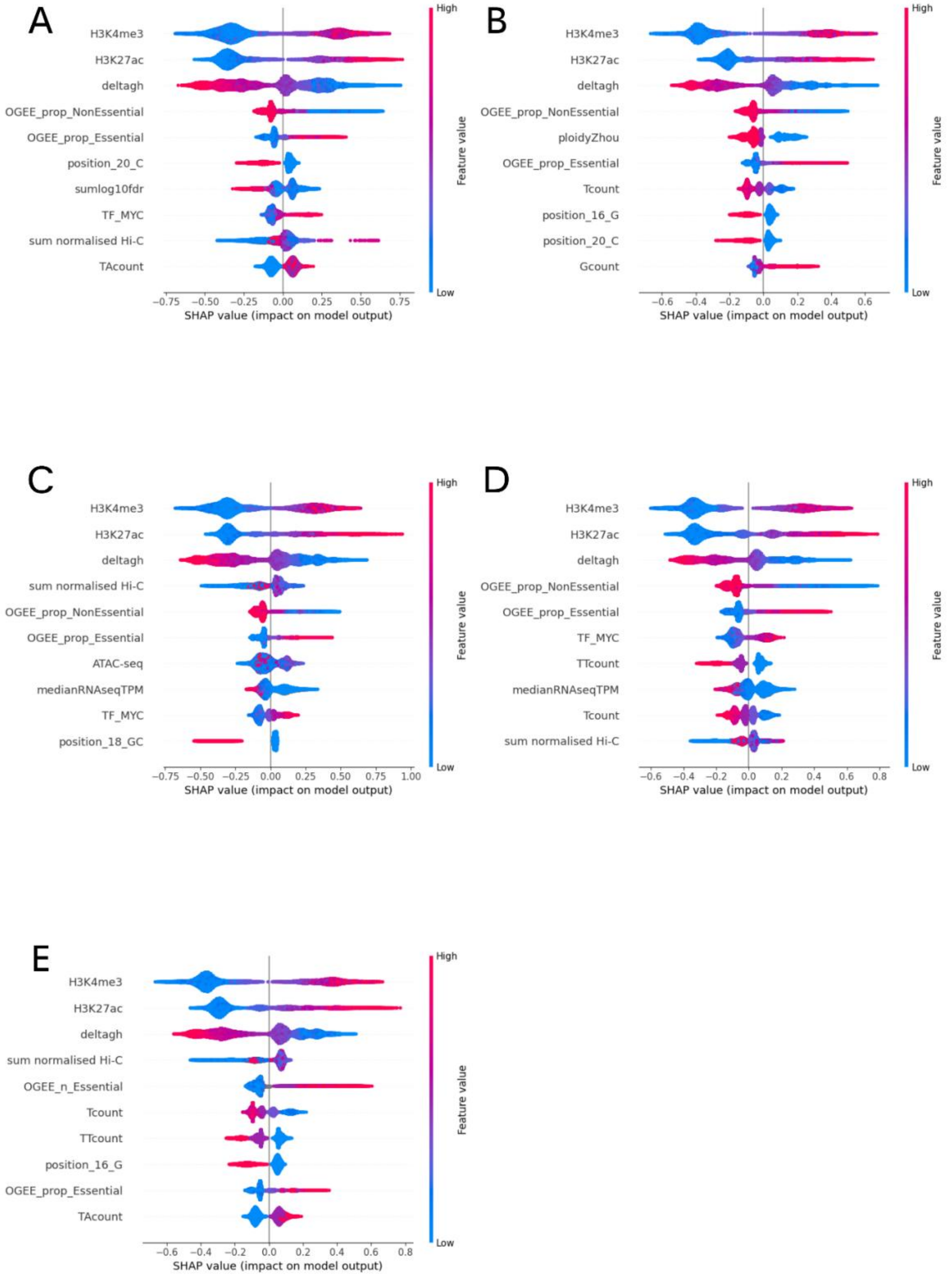


***Supplementary Fig. 2 | SHAP summary plot for gRNAs targeting promoter regions.*** *Top 10 significant features in the XGBoost predictive model for cell fitness, presented separately for each of the 5 folds of gRNAs.*

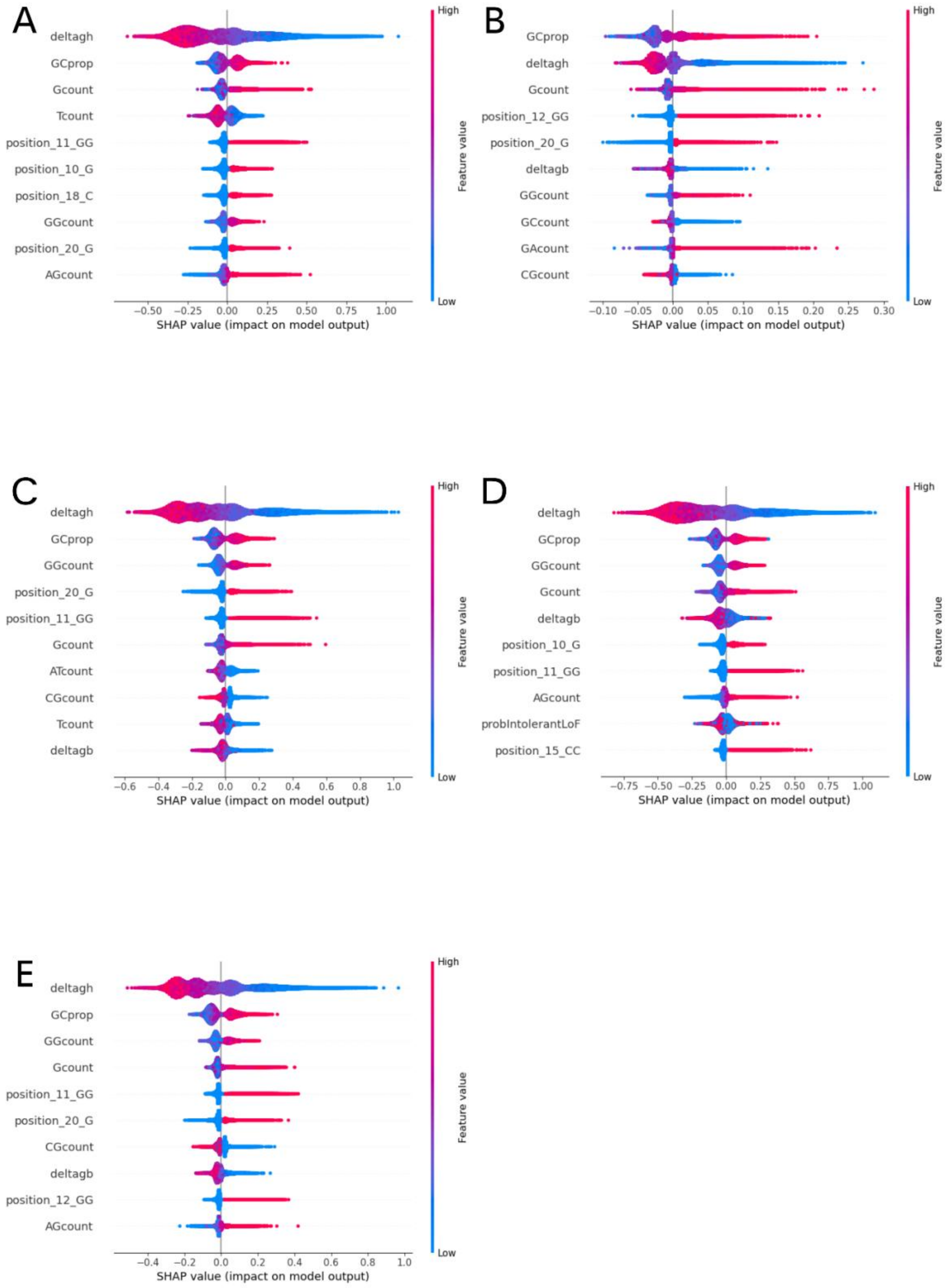


***Supplementary Fig. 3 | SHAP summary plot for gRNAs targeting enhancer regions.*** *Top 10 significant features in the XGBoost predictive model for cell fitness, presented separately for each of the 5 folds of gRNAs.*

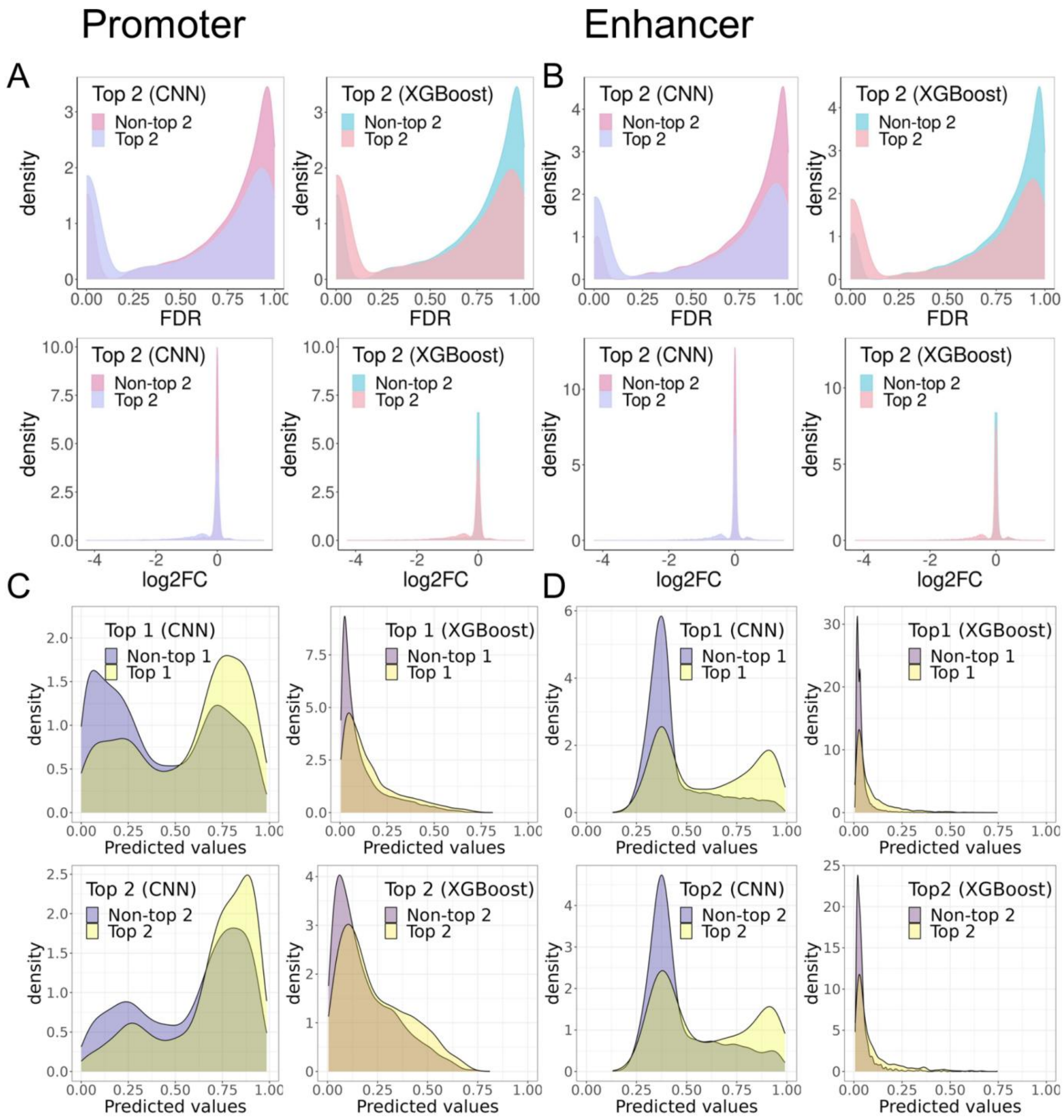


***Supplementary Fig. 4 |Predictive performance for cell fitness.*** *(A,B) Density plots of FDR and $log_2FC$ between the top 2 gRNA selected by CNN and XGBoost and the remaining gRNAs for promoter and enhancer, respectively. (C,D) Density plots of predicted probability of significance by CNN and XGBoost for the top-ranked observed gRNAs and remaining gRNAs in the promoter and enhancer, respectively.*

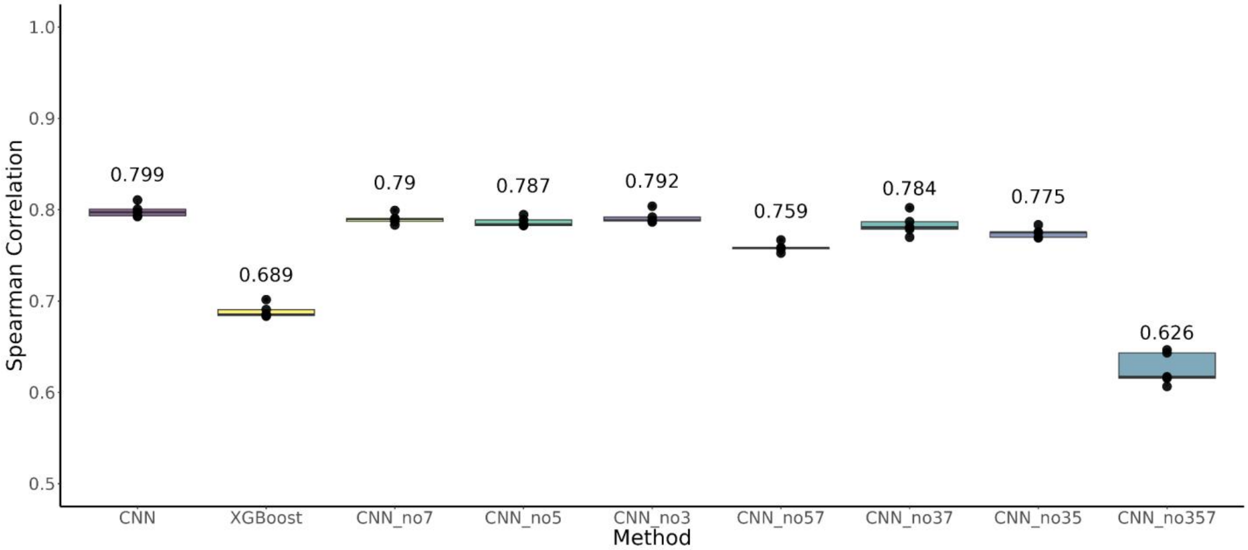


***Supplementary Fig. 5 | Prediction performance for counts in wild-type cells.*** *Mean Spearman correlation across five data folds are listed on top of each box. CNN_no57: CNN without kernels of size 5 and 7 nt; CNN_no357: CNN without kernels of size 3, 5, and 7 nt. P-values from paired t-tests are listed above the bars.*

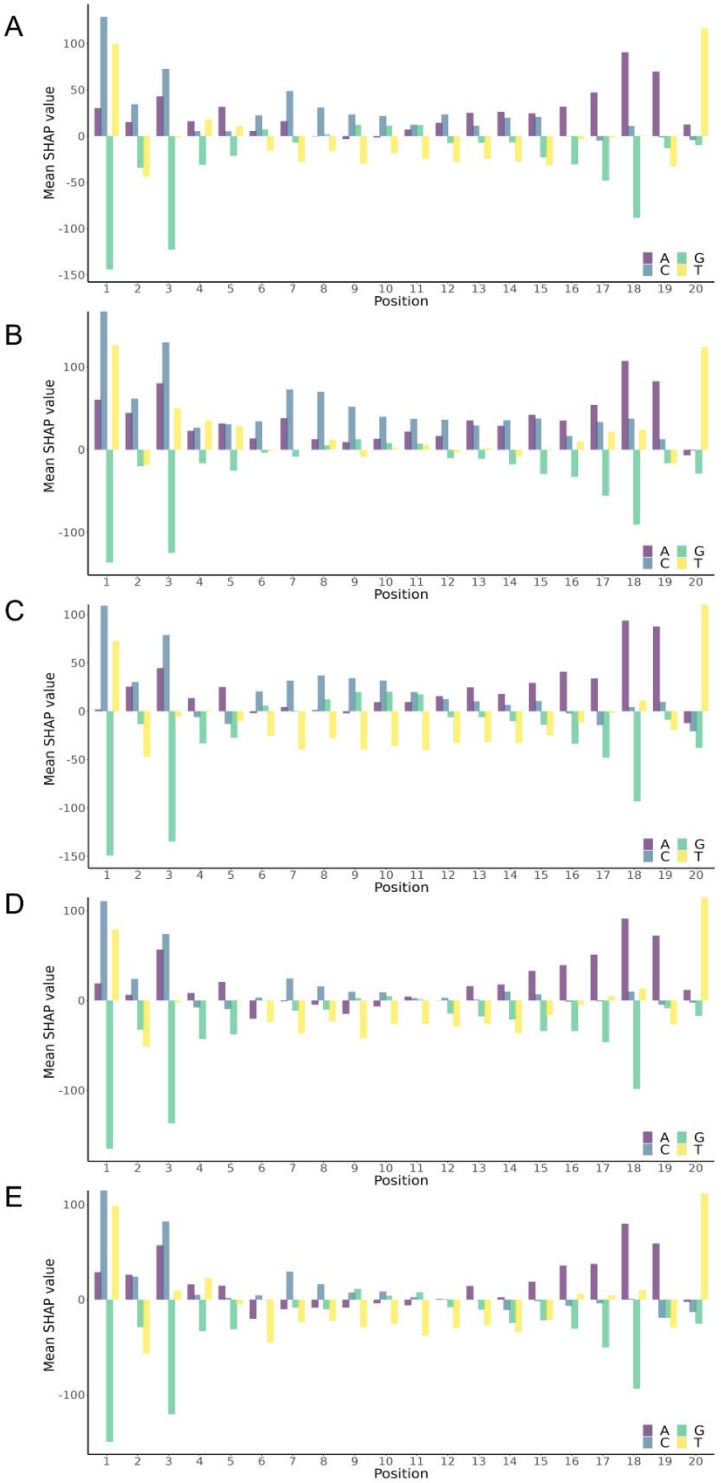


***Supplementary Fig. 6 | DeepSHAP summary plot.*** *Mean DeepSHAP values for input sequence positions in all 5 folds of gRNAs from trained wild-type count CNN models.*

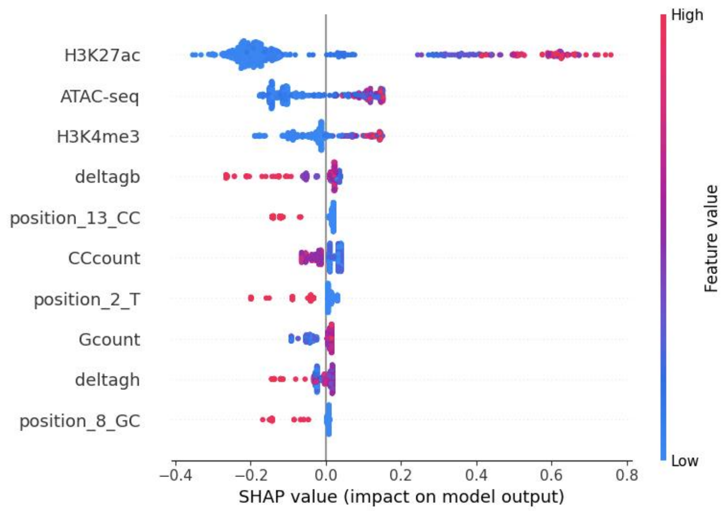


***Supplementary Fig. 7 | SHAP summary plot for scCERES.*** *SHAP summary plot that displays the top 10 significant features for XGBoost across all 5 folds of gRNAs.*

### Supplementary Table

##### Supplementary Table 1A: Sample size in different training folds for cell fitness data

| Task | Fold | Chromosomes Included in Test Sets | Significant gRNAs | Insignificant gRNAs |
|---|---|---|---|---|
| Cell Fitness - Promoter Regions | 1 | 4, 10, 16, 17, 21, X | 1,042 | 62,251 |
| | 2 | 1, 14, 22 | 775 | 47,273 |
| | 3 | 2, 7, 11, 13 | 864 | 50,450 |
| | 4 | 3, 5, 6, 18, 19 | 977 | 64,416 |
| | 5 | 8, 9, 12, 15, 20 | 815 | 51,217 |

| | | | | |
|---|---|---|---|---|
| Cell Fitness - Enhancer Regions | 1 | 4, 10, 16, 17, 21, X | 1,079 | 157,864 |
| | 2 | 1, 14, 22 | 853 | 122,124 |
| | 3 | 2, 7, 11, 13 | 984 | 143,593 |
| | 4 | 3, 5, 6, 18, 19 | 1,030 | 161,938 |
| | 5 | 8, 9, 12, 15, 20 | 899 | 143,187 |

**Supplementary Table 1B: Sample size in different training folds for wild-type cell count data**

| Task | Fold | Chromosomes Included in Test Sets | Number of gRNAs |
|---|---|---|---|
| Wild-type average counts | 1 | 16, 17, X | 1,968 |
| | 2 | 1, 14, 21, 22 | 1,975 |
| | 3 | 2, 4, 7, 11, 13 | 1,973 |
| | 4 | 3, 5, 6, 10, 18, 19 | 2,008 |
| | 5 | 8, 9, 12, 15, 20 | 1,951 |

**Supplementary Table 1C: Sample size in different training folds for single cell genome-wide screen**

| Task | Fold | Chromosomes Included in Test Sets | Significant gRNAs | Insignificant gRNAs |
|---|---|---|---|---|

| | | | | |
|---|---|---|---|---|
| Single cell genome-wide screen for K562 cells | 1 | 5, 6, 12, 15, 21, 22 | 107 | 450 |
| | 2 | 2, 4, 10, 16, 18 | 98 | 403 |
| | 3 | 1, 7, X | 104 | 437 |
| | 4 | 3, 8, 9, 14, 20 | 77 | 418 |
| | 5 | 11, 13, 17, 19 | 126 | 377 |

**Supplementary Table 1D: Sample size in different training folds for single cell MHC region**

| Task | Fold | Significant gRNAs | Insignificant gRNAs |
|---|---|---|---|
| Single cell screen in MHC region for K562 cells | 1 | 334 | 1,562 |
| | 2 | 308 | 1,599 |
| | 3 | 318 | 1,634 |
| | 4 | 380 | 1,567 |
| | 5 | 329 | 1,519 |
| Single cell screen in MHC region for NPC | 1 | 314 | 1,550 |
| | 2 | 252 | 1,618 |
| | 3 | 345 | 1,544 |
| | 4 | 387 | 1,531 |
| | 5 | 294 | 1,501 |

| | | | |
|---|---|---|---|
| Single cell screen in MHC region for iPSC | 1 | 519 | 1,374 |
| | 2 | 530 | 1,397 |
| | 3 | 513 | 1,364 |
| | 4 | 578 | 1,358 |
| | 5 | 524 | 1,413 |

**Supplementary Table 2A: Comparing gRNAs impact on cell fitness vs gRNAs impact in at least one gene expression**

| | gRNA significant in at least one gene expression | gRNA insignificant in any gene expression |
|---|---|---|
| gRNA significant in cell fitness | 295 | 586 |
| gRNA insignificant in cell fitness | 144 | 413 |

**Supplementary Table 2B: Comparing gRNAs impact on cell fitness vs gRNAs impact in larger than three gene expression**

| | gRNA significant in larger than three gene expression | gRNA insignificant in any gene expression |
|---|---|---|
| gRNA significant in cell fitness | 21 | 860 |
| gRNA insignificant in cell fitness | 2 | 555 |

#### Supplementary Table 3: ENCODE human functional annotations

| File name | Biological replicate | Type |
|---|---|---|
| **K562** | | |
| ENCFF512VEZ.bam | Rep1 | ATAC-seq |
| ENCFF987XOV.bam | Rep2 | ATAC-seq |
| ENCFF656DMV.bam | Rep1 | H3K4me3 |
| ENCFF440ARP.bam | Rep2 | H3K4me3 |
| ENCFF907MNY.bam | Rep1 | H3K27ac |
| ENCFF907MNY.bam | Rep2 | H3K27ac |
| ENCFF307XGY.bam | Rep1 | GATA2 |
| ENCFF032UMK.bam | Rep2 | GATA2 |
| ENCFF304UTB.bam | Rep1 | TAL1 |
| ENCFF631RZK.bam | Rep2 | TAL1 |
| ENCFF503ZCR.bam | Rep1 | MYC |
| ENCFF604CDP.bam | Rep2 | MYC |
| **WTC11 iPSC** | | |
| ENCFF235SBC.bam | Rep1 | ATAC-seq |
| ENCFF735XZN.bam | Rep2 | ATAC-seq |
| ENCFF979XKI.bam | Rep1 | H3K4me3 |
| ENCFF479NLV.bam | Rep2 | H3K4me3 |
| ENCFF738QRT.bam | Rep1 | H3K27ac |
| ENCFF696GDY.bam | Rep2 | H3K27ac |
| **NPC** | | |
| ENCFF589KAY.bam | Rep1 | H3K4me3 |
| ENCFF212FUS.bam | Rep2 | H3K4me3 |
| ENCFF805URT.bam | Rep1 | H3K27ac |

| | | |
|---|---|---|
| ENCFF120AVO.bam | Rep2 | H3K27ac |
| ENCFF477FHN.bam | Rep3 | H3K27ac |
